# A phase diagram for gene selection and disease classification

**DOI:** 10.1101/002360

**Authors:** Hong-Dong Li, Qing-Song Xu, Yi-Zeng Liang

## Abstract

Identifying a small subset of discriminate genes is important for predicting clinical outcomes and facilitating disease diagnosis. Based on the model population analysis framework, we present a method, called PHADIA, which is able to output a phase diagram displaying the predictive ability of each variable, which provides an intuitive way for selecting informative variables. Using two publicly available microarray datasets, it’s demonstrated that our method can selects a few informative genes and achieves significantly better or comparable classification accuracy compared to the reported results in the literature. The source codes are freely available at: www.libpls.net.

## Introduction

High throughput experiments such as DNA microarray allow quantifying the expression levels of thousands of genes simultaneously and provide a large amount of data of potential clinical use such as disease risk prediction and classification [8,9,12,14,22,31]. The goal of disease/cancer classification includes, but is not limited to, predicting prognosis, proposing therapy according to the clinical situation, advancing therapeutic studies *etc*. Thus, it is critical for physicians/clinicians to establish the rule for accurate classification of tumor before any treatment is administered to the patient in order to avoid unnecessary treatment or propose the most appropriate therapies. The study of gene expression data based cancer classification has been extensively reported [21,25,26,29,30,34]

Due to many factors such as high cost, expression data are usually measured for a large number of genes on a small number of samples, giving rise to a measurement data matrix of a few rows and many columns. Predictive modeling and variable selection for such data is known as the a “large *p*, small *n*” problem[4,10,35], which is very challenging and has received a lot of attention in bioinformatics and statistics. So far, many methods have been proposed to identify potential genes which are relevant to cancer classification, *e.g.*, t-scores, class distinction correlation[12], support vector machines (SVM) [11,24], Gaussian process[7], sparse logistic regression[6], regularized ROC method[22,23] entropy-based method [20], graph based feature classification[13] and the MIA approach [18]. In predictive modeling, it is our opinion that a variable or gene is predictive performance should be assessed in terms of two aspects: (1) the expectation and (2) variance of predictive performance of a variable. The former shows how a variable improves a model’s performance when included and the latter reflects how a variable affects the confidence interval (stability) of a model. These two factors are essential for quality assessment of variables but seemingly have not been addressed by existing methods.

In our previous work [18], we proposed the MIA (margin influence analysis) method that can computationally assess the expectation of predictive performance of variables using a population of sub-models [16,17,19]. However, the MIA method is only applicable to support vector machines [15] and does not consider the prediction variance of variables. Based on model population analysis [16,17,19], here we report a computational approach which can output a phase diagram displaying quality of variables in a two-dimensional plot. The algorithm was termed PHADIA. The phase diagram provides an intuitive tool for variable selection. Though our algorithm is implemented with PLS-LDA (Partial least squares - linear discriminant analysis), the PHADIA can be applied in combination with any other classifiers such as support vector machines and Bayesian network classifier. We applied PHADIA to two publicly available gene datasets and evaluated its performance for gene selection.

## Methods

As mentioned above, PHADIA is proposed based on MPA[5,16], which is a general framework for developing new methods by analyzing the interesting model-related parameters, *e.g.* prediction errors and variable coefficients, of a number of sub-models based on Monte Carlo Sampling (MCS). An MPA algorithm consists mainly of three steps: (1) sampling N sub-datasets randomly, (2) building a sub-model using each sub-dataset, and (3) statistically analyzing some interesting output, *e.g.* prediction errors, of all the N sub-models. The third step is the key point of MPA. In the following section, the PHADIA algorithm will be described according to these three steps.

### Sub-dataset sampling in the variable space

Given a dataset (**X**, **y**), where **X** of size n×p has n samples and p variables and **y** of size n×1 records the class label of each sample, valued 1 or −1 in the binary classification situation. The sub-dataset sampling in the variable space is described in the following (Figure 1). The number of Monte Carlo sampling (MCS) is set to N (*e.g.* 10,000). At each sampling, Q out of the p variables will be randomly sampled, giving rise to a sub-dataset of size n×Q. Repeating this procedure for N times, N sub-datasets in total can be obtained, which are denoted (**X**_sub_, **y**_sub_)_i_, i = 1, 2, 3, …N. In Figure 1, the filled squares stand for the sampled variables.

**Figure 1.**
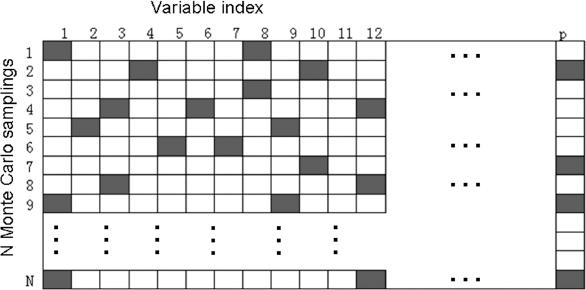
Illustration of Monte Carlo sampling in the variable space. In each sampling, a sub-dataset containing only a given number (Q, e.g. 10) of variables are randomly selected from the original data, denoted by filled square. Finally, N sub-datasets are generated and a sub-model will be built using each of the sub-dataset.

### Sub-model building using PLS-LDA

For each sub-dataset, a PLS-LDA model is built with the number of latent variables optimized by cross validation [2,32]. The reason to choose PLS-LDA is that (1) PLS has the potential to deal with high dimensional data and (2) linear classifier is easy to interpret compared to the nonlinear methods. N PLS-LDA classifiers are established in this step. 5-fold cross validation[3,28] is used to assess the performance of each sub-model. So each model is associated with a prediction error.

### Statistical analysis of prediction errors for computing phase diagram

Without loss of generality, we take the *i*th variable for example to illustrate the procedure for computing a phase diagram.

First, we partition all the previously computed N PLS-LDA models into two groups, say Group A and Group B. Group A collects all the models including the *i*th variable. The remaining models not including the *i*th variable belong to Group B. Assuming that the numbers of the models in Group A and B are N_A_ and N_B,_ respectively, the sum of N_A_ and N_B_ is equal to N. So we have N_A_ prediction errors for Group A and N_B_ prediction errors for Group B. Then, the mean and standard deviation of the N_A_ and N_B_ prediction errors can hence be easily calculated, denoted MEAN_A_, SD_A_, MEAN_B_ and SD_B_, respectively. Finally, two statistics are can be computed for assessing the *i*th variable’s prediction ability. The first is defined as the difference between MEAN_A_ and MEAN_B,_ which is denoted and computed using the following formulae.

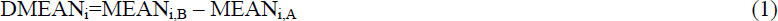

Clearly, DMEAN_i_ measures the increment of prediction performances of model including the *i*th variable over models without it. So, if DMEAN_i_>0, one may infer that the prediction ability could be improved if a model contains the *i*th variable, and vice versa. To judge whether the mean errors of Group A and Group B are different, the nonparametric Mann-Whitney U test is employed to calculate a p-value which in combination with DMEAN is able to tell whether a variable can significantly improve prediction performance or not. In analogy to DMEAN, another statistic is defined as:

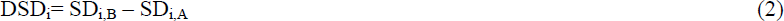

By definition, DSD_i_ can be thought as an criterion describing whether the inclusion of the *i*th variable will increase the variance of a model or not. It could be expected that the *i*th variable has the potential to stabilize a model if DSD_i_>0.

After computing DMEAN and DSD for each variable, one can plot DSD against DMEAN for all the variables. Such a plot is called a phase diagram in our work which intuitively displays the quality of all variables together. The phase diagram is illustrated in Figure 2. In this figure, all the variables are grouped into four regions. Phase 1: DMEAN>0, DSD>0, containing informative variables that can reduce prediction errors and also reduce prediction variance; Phase 2: DMEAN<0, DSD>0, housing variables which can will increase prediction variances; Phase 3: DMEAN<0, DSD<0, corresponding to variables that decrease model performance but can reduce prediction variances; Phase 4: DMEAN>0, DSD<0, including the “worst” variables which not only increase prediction errors but also increase prediction variances. Based on p-value of each variable, the shadow region with DMEAN close to 0 indicates variables which will not have significant influence on a model’s prediction error. In general, variables in Phase 1 are best performing and those in Phase 2 can also be considered to be included into a model. However, the variables in Phase 3 and 4 are not suggested to be used for building a model.

**Figure 2.**
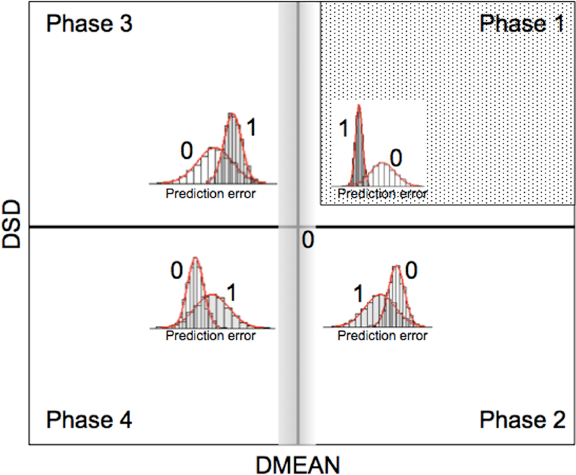
A schematic of the phase diagram for variable selection. Based on the DMEAN and DSD values (see main text for definition), variables fall into four main phases. The variables in Phase 1 can not only increase a model’s performance (DMEAN>0) but also reduce prediction variances (DSD>0). In the inset, the peak denoted by “1” stands for the prediction error distribution of models including such a variable, while the peak denoted by “0” is the prediction error distribution of models that do not include this variable. This type of variables is called informative variables. The variables in Phase 2 can also increase performance but at the cost of increased variance, thus being less informative than those in Phase 1. In contrast, variables in Phase 3 and 4 will reduce the performance (DMEAN<0) of models, so they are suggested not to be used for modeling. These are called interfering variables. Specifically, including variables in Phase 4 into a model will even increase the prediction variance (DSD<0). In addition, variables at the boundary (the shadow region with DMEAN close to 0) can not significantly increase or decrease model performances, and are hence thought of as being uninformative.

## Results and discussion

### Colon data

This dataset contains the expression values of 6500 human genes measured on 40 tumor and 22 normal colon tissues using the Affymetrix gene chip. 2000 genes with the highest minimal intensity across samples were selected by Alon *et al.* [1]. Gene expression values were log2 transformed before modeling.

The parameter N, the number of Monte Carlo samplings, for PHADIA is fixed 10,000. Four Q values [20, 50, 100, 200] were tested. For each Q, we ran the PHADIA algorithm 10 times and calculated the predictive performance of PLS-LDA using cross validation (Figure S1). Considering both the mean and variance of prediction errors, Q=20 was chosen to be the optimal one. Although the sub-dataset is chosen randomly, we found that the top ranked informative variables identified by the PHADIA algorithm is highly reproducible (**Table S1**).

Figure 3 shows the phase diagram from one run of the PHADIA algorithm. This plot is divided into four phases, corresponding to the four types of genes as described previously. The genes (probe sets) in Phase 1 are most informative. Phase 2 houses less informative genes which can still be considered to be included in model. However, those genes in Phase 3 and 4 will most likely reduce a model’s performance and therefore should be eliminated as interfering genes. We found that there are 878 genes with DMAN>0 (in Phase 1 or Phase 2), out of which 206 are significant with a p-value < 0.05.

**Figure 3.**
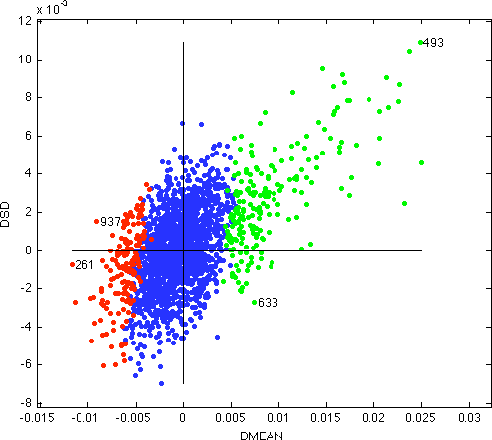
The phase diagram of the colon data. Red: informative variables (DMEAN>0, p<0.05); blue: uninformative variables (p>0.05); red: interfering variables (DMEAN<0, p<0.05).

We then from each Phase, manually picked one gene (Phase 1: index=493, Phase 2: index=633, Region 3: index=937 and Phase 4: index=261) and displayed their prediction error distributions in Figure 4. Take the 493th gene for example, the prediction error distribution of models including this variable shows significantly lower mean errors and smaller variance, representing the most informative variable. In contrast, the variables in Plot A and C are of poor predictive performances since including them into a model will on average reduce a model’s performance.

**Figure 4.**
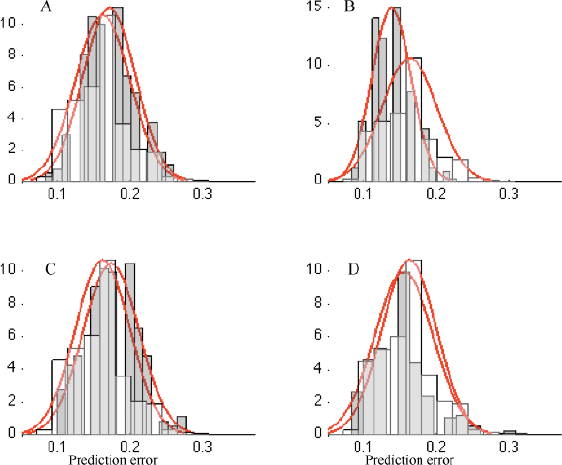
The prediction error distributions of four genes from each of the four phases for the colon data. They are picked from Phase 1 (plot B, index=493), Phase 2 (plot D, index=633), Phase 3 (plot A, index=937) and Phase 4 (plot C, index=261) in Figure 3, respectively.

To build a parsimonious PLS-LDA classifier, we ranked all the variables based on their DMEAN value and selected a subset of 200 genes using a forward strategy. The achieved average median leave-one-out-cross-validation (LOOCV) error of 10 replicate runs of the PHADIA algorithms is 0.0806±0.0221. For the same data, Furey *et al.* (2000) misclassified 6 samples using SVM based on LOOCV, leading to a prediction error = 0.097. In Nguyen *et al.* (2002), their best result is obtained using PLS including 50 or 100 genes with the misclassification error=0.065, which are slightly better than ours. Compared to the reported results, the proposed PHADIA algorithm shows competitive performances.

### Estrogen data

This dataset consisted of the expression values of 7129 genes of 49 breast tumor samples, and presented by West *et al.* (2001) and Spang *et al*.(2001)[27]. There are 25 LN+ samples and 24 LN-samples. Before gene selection and classifier building, pretreatment is done on this data following the same methods described in Ma *et al.* (2005). 3333 genes were left and log2 transformed in our analysis.

For this data, N is also set to 10,000, and the optimal Q value is determined to be 20 using cross validation (Figure S2). The results of PHADIA are shown to be highly reproducible (**Table S1**). The phase diagram from one run of PHADIA with Q=20 is shown in Figure 5. In this plot, the 77th gene stands out, which should be of high predictive value based on our method. For illustration, we also selected one gene from each phase and their prediction error distributions are shown in Figure 6. Take the 77th gene as an example, it remarkably reduces the prediction error while simultaneously improve the predictive stability in terms of lower variance if included in a model. It may be inferred that the 77th gene is a key factor for underling the physiological state of estrogen. From Phase 1 and Phase 2, 312 genes were found to be informative with p<0.05.

**Figure 5.**
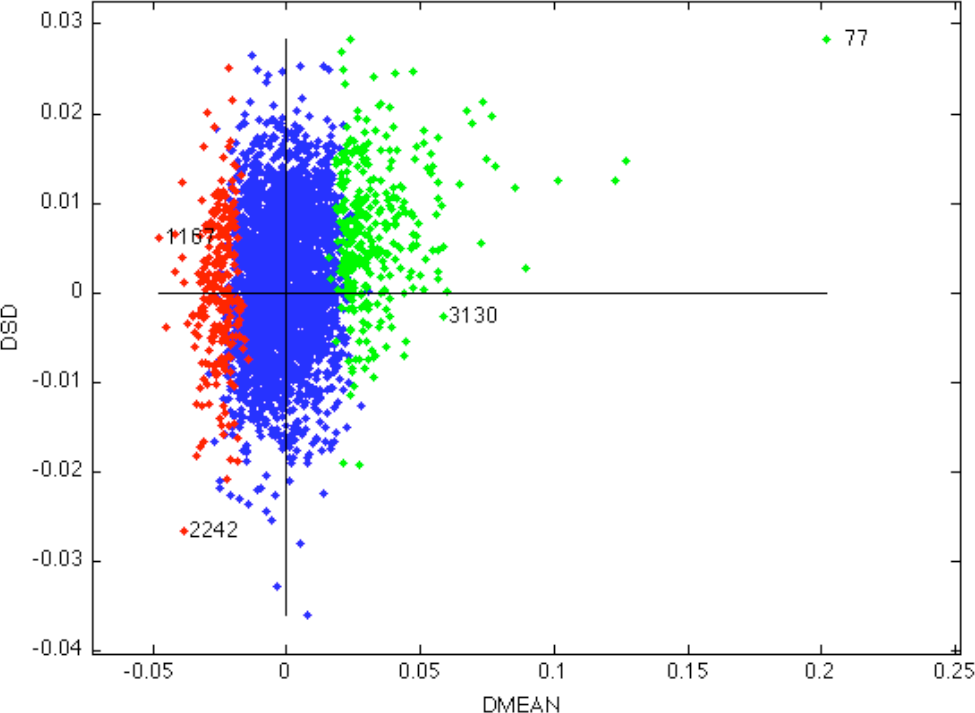
The phase diagram of the estrogen data. Red: informative variables (DMEAN>0, p<0.05); blue: uninformative variables (p>0.05); red: interfering variables (DMEAN<0, p<0.05).

**Figure 6.**
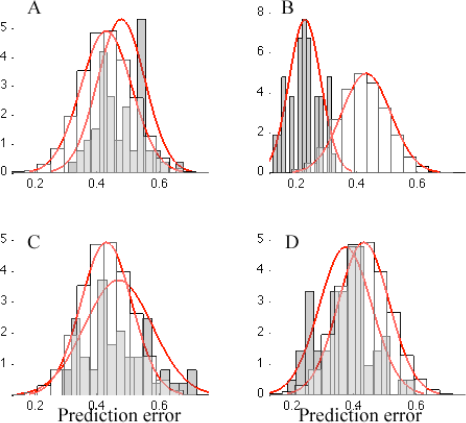
The prediction error distributions of four genes from each of the four phases for the estrogen data. They are picked from Phase 1 (plot B, index=77), Phase 2 (plot D, index=3130), Phase 3 (plot A, index=1167) and Phase 4 (plot C, index=2242) in Figure 5, respectively.

Using the same method as applied to the colon data, we selected 25 genes and built a PLS-LDA classifier with a median LOOCV error 0.06±0.00 over 10 replicate runs of the PHADIA algorithm. For this dataset, based on their selected 100 genes, Dettling and Buhlmann (2003) yields classification errors 0.020 (LogitBoost, optimal), 0.06 (AdaBoost, 100 iterations) and 0.040 (CART). In the work of Ma *et al* (2005), their reported misclassification errors are 0.120±0.080. These results demonstrated that our method is very compelling.

## Conclusions

Based on model population analysis, we in the present study introduced the PHADIA algorithm for variable selection. One unique feature of PHADIA is that it can output a phase diagram that describes the prediction ability of variables in terms of the expectation and variance of their prediction errors if included in a model. Based on the phase diagram, variables can also be classifier into informative, uninformative and interfering ones in a similar manner as in our previous work [18,19,33]. When applied to two gene expression datasets, competitive performances were achieved compared to the results reported in the literature. Our results indicate that PHADIA algorithm is a powerful tool for visualizing variables’ prediction ability and identifying informative variables. It’s expected that PHADIA will find more applications in other fields.

## Acknowledgements

This work is financially supported by the National Nature Foundation Committee of P.R. China (Grants No. 21075138). The studies meet with the approval of the university’s review board.

## Supplementary materials

**Figure S1.**
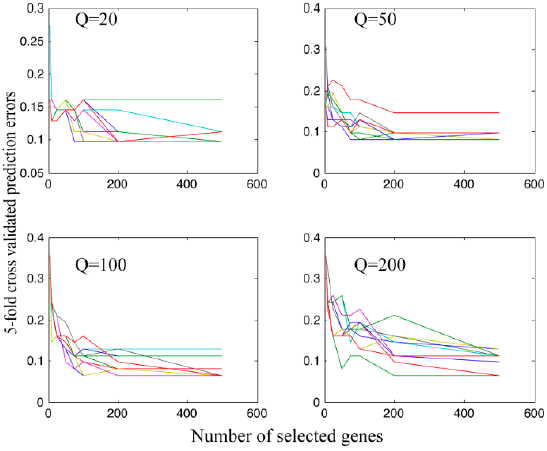
5-fold cross validated prediction errors using different Q values for the colon data.

**Figure S2.**
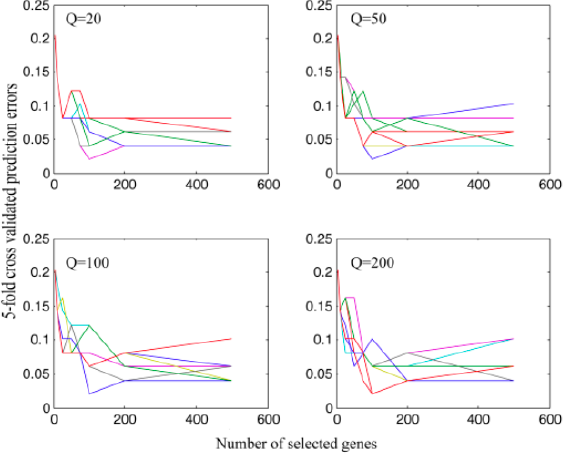
5-fold cross validated prediction errors using different Q values for the estrogen data.

